# Molecules to Mankind: Bridging Laboratory and Population Training

**DOI:** 10.1101/821587

**Authors:** Rachel M. Burke, Julie A. Gazmararian, Nael A. McCarty, Benjamin L. Rambo-Martin, Kelly A. Shaw

**Author notes:** Corresponding Author: Julie Gazmararian, PhD, MPH, Professor, Department of Epidemiology, Rollins School of Public Health, Emory University, 1518 Clifton Road, Atlanta, Georgia 30330.

## Abstract

**Aim/Purpose:** Today’s biomedical researchers are expected to apply understanding of basic biology to improve human health. Meeting this goal requires mastery of both laboratory and population sciences, each of which has its own knowledge base, techniques, and training paradigms.

**Background:** Emory University’s “Molecules-to-Mankind” (M2M) doctoral pathway was established in 2009 to be an evolving model of interdisciplinary PhD education. M2M supplements fellows’ home programs, ensuring they receive training in both population and laboratory sciences.

**Methodology:** The present paper describes the M2M program in detail. Surveys of faculty and fellows were also carried out, and the results are presented herein.

**Contribution:** The M2M program follows a unique model by which doctoral students receive training in both population and laboratory sciences. The present paper describes this model, such that the information can be disseminated to other educational institutions interested in implementing similar programs.

**Findings:** This unique model facilitates engagement of stakeholders including the fellow’s home program, dissertation advisor, and Emory’s professional schools. Recruited across biomedical PhD and MD/PhD programs, fellows have diverse research experiences and represent “spokes” bound together by the M2M “hub.” This hub’s central feature is a weekly seminar class where fellows and faculty members gather for open discussion with interdisciplinary speakers with successful research careers, emphasizing speakers who have tied laboratory and population sciences in their own work. This forum provides an encouraging environment for dialogue on all aspects of biomedical research from the science itself, to the speaker’s career path, and the logistics of garnering institutional support and building transdisciplinary collaborations. A decade since its inception, M2M has a proven track record of shaping early-stage careers for its 49 alumni to date.

**Recommendations for Practitioners:** Engaging trainees outside their home doctoral programs can have positive implications on overall quality of education.

**Recommendation for Researchers:** As interdisciplinary models grow ever more important in biomedical sciences, it is important to further examine the best teaching methods for training the next generation of scientists.

**Impact on Society:** Interdisciplinary training at the doctoral level is important to produce future cohorts of engaged and versatile scientific leaders.

**Future Research:** Future research should continue to explore novel methods of training graduate students at the doctoral level.

## INTRODUCTION

Historically, scientific education at the doctoral level has focused on developing specialists—professionals with a deep but narrow band of knowledge. Laboratory scientists often train extensively in molecular subjects but are not exposed to training in population health disciplines such as epidemiology or demography. Meanwhile, most population scientists do not receive training in laboratory science topics. Yet the need for interdisciplinary research grows ever greater, as technological advancements increase our ability to generate more and “bigger” data, and as our research questions grow ever more complicated and systems-based. In today’s landscape, scientists must be prepared to relate their research to broader questions, to articulate policy implications, and to communicate to diverse scientific as well as non-scientific audiences. Scientists trained in both population-and laboratory-based sciences are uniquely equipped to face these challenges and to contribute to interdisciplinary research at a high level.

Recognizing the need to educate this new cadre of scientists, the Burroughs Wellcome Fund (BWF) released an initial call for proposals in 2008 for the Institutional Program Unifying Population and Laboratory Based Sciences (PUP) award. The goal of this award was to establish a program that would build upon the strengths of predoctoral training in specific fields emphasized at each institution in order to train a cohort of researchers who can bridge the gap between population-and laboratory-based sciences. Programs were expected to experiment with models of training to find what worked at their institution, and then incorporate those effective interventions into the fabric of graduate training for the long term. In 2009, three universities, including Emory University in Atlanta, Georgia, received the inaugural round of the training grant, providing $500,000 annually for five years. In the fall of 2010, the first cohort of Emory students was enrolled into the “Human Health: Molecules-to-Mankind” (M2M) program. M2M was designed not as a separate PhD program, but as a “doctoral pathway” within the Laney Graduate School (LGS). Students were required to first be accepted into one of Emory’s PhD programs as their “home program,” and then apply separately to M2M before matriculating into that home program. Some students entered M2M from a population sciences background, while others entered from the perspective of laboratory sciences. The M2M curriculum and requirements are complementary to, but separate from, a student’s home program requirements, and are designed to help students bridge the gap between population-and laboratory-based sciences.

## Description of M2M

### Program Structure

The original vision for the M2M Program espoused a doctoral pathway with the theme “Understanding human health: integrating biology, behavior, environments, and populations.” The program creators suggested that Emory University was an ideal institution for investment in an interdisciplinary program to enhance graduate education tying together population and laboratory sciences. Emory University is geographically compact and walkable between academic units (distances mostly 10-15 minutes by foot), with LGS graduate faculty housed across diverse schools, such as those composing the academic side of the health sciences center (Medicine, Nursing, Public Health), the schools of Law, Theology, and Business, as well as the Emory College of Arts and Sciences. Many LGS faculty have clinical responsibilities in addition to their teaching and research, facilitated by the proximity of Emory University Hospital and The Emory Clinic, located on the Emory campus. The structure of a central graduate school (LGS) and the co-location of the other academic units promotes interdisciplinary thought, discussions, pedagogy, and research. Furthermore, Emory is physically contiguous to one of three Children’s Healthcare of Atlanta hospital locations as well as the Centers for Disease Control and Prevention (CDC). The national headquarters of the American Cancer Society, The Carter Center, and the headquarters of the international relief organization CARE are also located within six miles of Emory’s campus. Emory also partners with the Georgia Institute of Technology (Georgia Tech), only five miles away.

Facilitated by Emory’s geographical compactness, the LGS cuts across all academic units and enables seamless registration of PhD students in any program. The LGS identifies and appoints the graduate faculty. Governance by that faculty is responsible for policies, program quality, curriculum review, and establishment of best practices. This arrangement greatly reduces barriers that might otherwise arise in the establishment of interdisciplinary, interdepartmental programs as envisioned for M2M. Over 200 new trainees begin life-sciences-related PhD programs at Emory every year. Since 2010, 49 students (ranging from 4–8 per year) have joined M2M from 11 of these programs (Table 1).

**Table 1.**
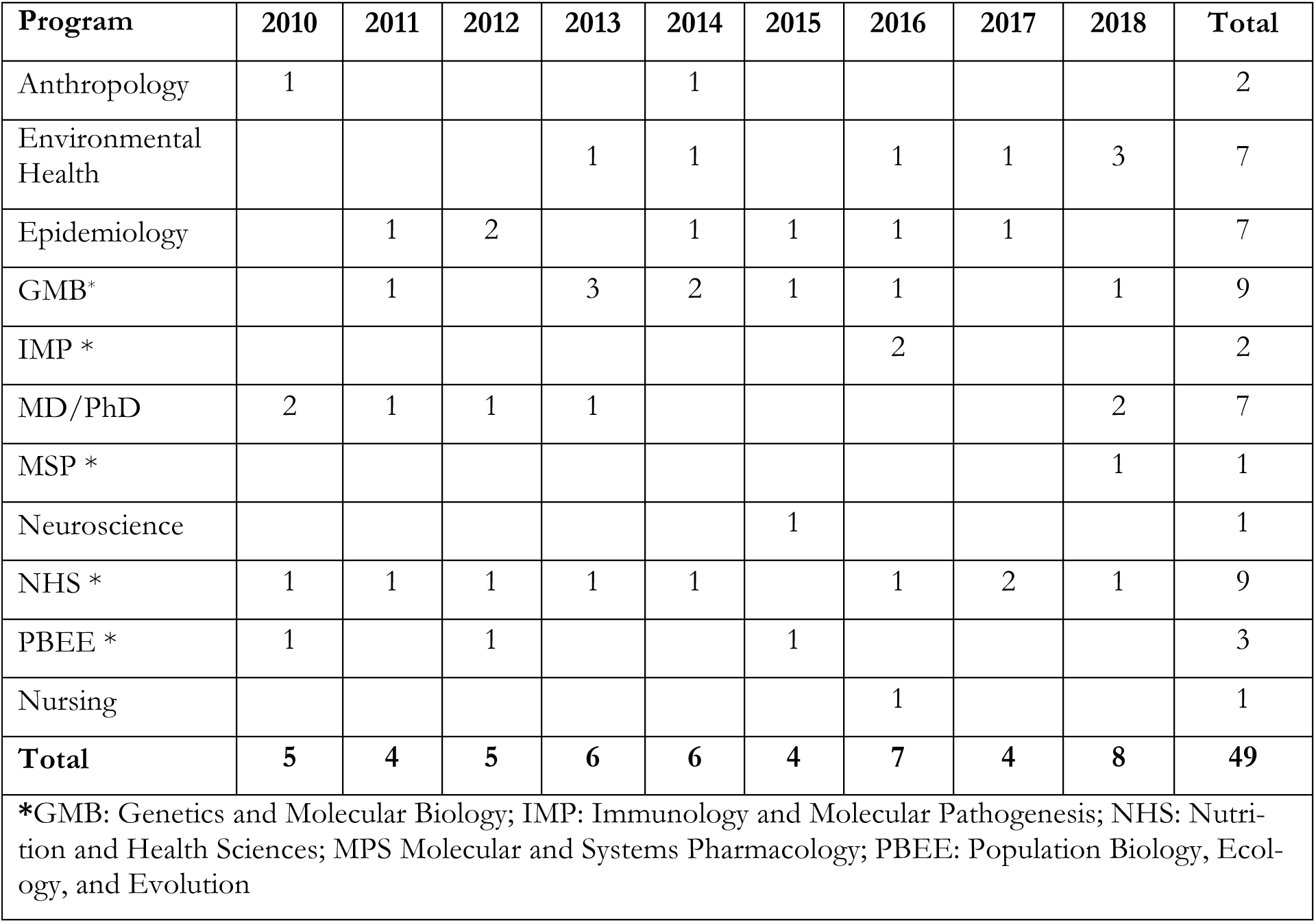
Home programs represented by M2M fellows, 2010-2018, n=49

With the goal of integrating laboratory and population sciences, M2M included four initial “tracks” that lend themselves especially to transdisciplinary education and for which there already existed established commitments and academic strengths at Emory. These tracks included: (1) Predictive Health; (2) Population Processes and Dynamics of Infectious Diseases; (3) Biomarkers and the Development of Acute and Chronic Diseases; and (4) Public Health Genomics: Genetic and Environmental Determinates of Health. Student applicants were asked to specify a track of interest, although these were not limiting in any way.

Each track was originally developed around the expertise of several faculty members with the appropriate background, and directed by two faculty track leaders: typically one that approached the track from the population perspective and one that approached it from the laboratory science perspective. Other faculty from across Emory’s various schools were involved in generating the proposal and became the initial program faculty. After the program began, other faculty with broad expertise and an interdisciplinary mindset joined upon expressing interest; mentors of program students often joined as program faculty.

### Recruitment

When M2M was first created, graduate students entered the program at the same time they entered their home program. However, difficulty coordinating with home programs during recruitment led to reduced support of M2M by the leaders of those home programs. Furthermore, students needed to be able to focus on orientation into their home program and their first-year courses before taking on M2M as additional training. As a result of these challenges, the model was changed such that students apply for entrance later, usually in their 2^nd^ or 3^rd^ year. Students further along in graduate school also are admitted to the program, but the focus of recruitment is on earlier years in order to emphasize interdisciplinary training throughout graduate school, and so that involvement in M2M could impact the student’s dissertation research to accomplish the goal of bridging disciplines. Information sessions and listserv announcements are used to promote recruitment, but the best recruiting tool is often word of mouth from current M2M students, who are the most outspoken advocates for peers applying to the program.

A major focus of M2M has been to expose students to hands-on training in both laboratory and population research. This was easy when students began M2M and rotations at the same time, but most 2^nd^ and 3^rd^ year students entering M2M have already completed rotations and chosen primary advisors. The program still seeks to ensure that students either had a broad exposure to both “wet lab” molecular and “dry lab” population research during their rotations, or have at least one experience in the other realm of research while working in their primary field after joining a lab group. M2M also requires students to have a second advisor that complements the background of the primary advisor, to help ensure bridging.

In its original implementation, M2M admitted students to the program for five years. However, as the program evolved this was limited to two years of official enrollment, although many students continued to attend program-related classes and activities well after their official enrollment ended.

### Program Requirements

There are minimal requirements added to students’ academic programs; M2M faculty track leaders help students identify coursework that will bolster their educational needs to cross the population-laboratory divide. Because M2M students are from very diverse programs and backgrounds, basic statistics and epidemiology coursework are required to get everyone on the same page. In addition, students are required to attend the M2M seminar class each semester they are officially enrolled in the program. The seminar class features speakers from diverse backgrounds and fields of study and also offers the opportunity for M2M students to present their own work to a supportive and constructive multi-disciplinary audience of faculty and students. The seminar class was designed to be a dynamic, discussion-driven experience each week rather than a typical academic presentation. Early on there was some difficulty with speakers who felt more comfortable with the usual seminar talk format, but this was resolved by emphasizing the goal of interactive discussions in speaker invitations. Speakers were explicitly asked to share personal stories of their career journey, especially how they tied lab and population sciences together. M2M values student input and has fostered a culture that promotes student engagement and participation in the course. Students often cite this course as the most valuable aspect of M2M, not only because of the groundbreaking work presented, but also the opportunity to have an actual dialogue with the speaker and their fellow classmates, who offer diverse perspectives and approaches to topics. Compared to traditional seminar classes in the home programs, which tend to be highly specialized, the diversity of approaches to science that are described by speakers in the M2M seminar class serves to greatly broaden the exposures that M2M students receive during their training.

Professional development also has become more of a focus in recent years, as many students seek to explore careers outside academia – a decision to which some faculty outside of M2M are openly hostile. To facilitate this exploration, many of the speakers presenting in the M2M seminar class are from outside of academia, and all are invited to give a brief background on what led them to their current position. In this way, students are exposed to the diversity of career paths available in industry, government, and non-profit sectors, as well as academia. There are also class sessions dedicated entirely to career development, and a description of resources available to the students. Examples of such sessions include a focus on what lies behind the curtain in academic positions (management of lab funds, service contributions, institutional politics, etc.) and how to choose a good academic institution for a career. With the recent emphasis among federal funding agencies on re-tooling graduate education, exemplified by the National Institute of General Medical Sciences and their emphasis on “Catalyzing the Modernization of Graduate Biomedical Training,” M2M sessions also include discussions of operational and professional skills including project management, leadership, self-awareness, scientific identity, team science, inclusivity, teaching, and communicating science. Though these skills are vital in modern research, they are missing from the curricula of many graduate programs.

In addition to the valuable experience of interacting with peers, M2M also offers unique support and participation from professors involved in the program. M2M provides an opportunity to engage with research faculty in open and honest conversation, removed from the politics of the student’s program or committee. The professors attending the seminar class are not grading the M2M fellows; rather, they are engaging as colleagues and mentors. Involvement of faculty dedicated to forward-thinking approaches to interdisciplinary training creates a space where students feel validated, rather than judged, for any career aspirations outside academia.

Other key features of M2M training include site visits to broaden intellectual horizons, such as learning about health disparities at the CDC Museum, environmental sustainability at Emory’s WaterHub water reclamation plant, and veteran engagement in research through the Atlanta Veterans Affairs Hospital. In addition, every year students and faculty nominate popular science books that are relevant to the broad concerns of the program; books are purchased for all program students and faculty, including the students accepted into the program for the upcoming fall semester, and treated as a summer reading assignment. Past readings have included *The Immortal Life of Henrietta Lacks* (Skloot), *The Emperor of All Maladies* (Mukherjee), and *The Drunkard’s Walk: How Randomness Rules Our Lives* (Mlodinow). When class reconvenes after summer, there is a student-led discussion about the book at the first M2M class session.

Lastly, there is an annual retreat held every spring. This retreat is an opportunity for students to provide updates on their research, professional development, and accomplishment of various graduate school milestones. Often, students describe challenges or roadblocks they are facing, and other students or faculty offer suggestions or assistance to help students continue to move forward. This meeting also serves as a forum to discuss M2M’s policies, with trainee and mentor voices given equal value.

### Additional Benefits to Students

All M2M students have access to professional development funds (up to $5,000 per year per student) that allow them to pursue opportunities that their graduate programs or mentors cannot or will not fund (e.g., workshops to learn research skills or laboratory supplies to support student work not directly related to their mentor’s focus). In the original model under which M2M operated, full stipend support was available for the first five years of a student’s engagement in the program. However, to enable the program to engage a larger number of students, and thereby increase the program’s impact, M2M has undergone structural changes that included decreasing stipend support and reducing the length of time that each student was in the program. In the first iteration of change, students were accepted into the program for a two-year period, and received only half stipend; professional development funds were held constant. In the second iteration, leading to the current state, students no longer receive stipend, but professional development funding remained the same. As stipend support decreased, the scope of program requirements decreased proportionally. As a result, the Program in its present iteration operates more like an NIH T32-funded training grant program (although without tuition support). Interestingly, these changes that included the phasing-out of stipends have not affected applications to the program, which indicates that the interest in this unique multi-disciplinary training, the seminar class, community of fellow students and faculty, and professional development provide tangible non-monetary benefits unmatched by other opportunities in graduate school, including those of the home programs.

## LITERATURE REVIEW

There has been increasing recognition that interdisciplinary approaches are necessary to tackle today’s complex research challenges[1,2] (often requiring data science in addition to another scientific domain[3]), and interdisciplinary collaborations have been on the rise both within and between institutes of higher learning[4]. Interdisciplinary research has also been championed by two of the largest sources of scientific funding in the United States: the National Institutes of Health and the National Science Foundation[5,6]. At the same time, several groups have noted the vital importance of interdisciplinary training during graduate education[7-9]. While there have been increases in interdisciplinary teaching and student degree offerings[4], including development[10,11], and implementation[12] of recommended core competencies, one researcher contends, “universities have tended to approach interdisciplinarity as a trend rather than a real transition and to thus undertake their interdisciplinary efforts in a piecemeal, incoherent, catch-as-catch-can fashion rather than approaching them as comprehensive, root-and-branch reforms”[13]. Programs such as those funded by the National Science Foundation’s Integrative Graduate Education and Research Traineeship have been developed in an attempt to fill in gaps and provide students with more structure and support for interdisciplinary training[14-17]. However, challenges to interdisciplinarity persist, including funding and forging successful partnerships[1] and the existing structure of doctoral programs[18]. Lessons learned from other programs such as Molecules to Mankind at Emory could be valuable to successful development of structured interdisciplinary training programs elsewhere.

## METHOD

A survey was developed to assess student and alumni attitudes towards the M2M program. Questions were open-ended, and covered the following themes: initial interest in M2M, impact of M2M on overall graduate student experiences, strengths and weaknesses of the program, and views on interdisciplinary training. This questionnaire was emailed out to all current students (11) and alumni (33), to be filled in via Google Sheets. The survey was open and functional for one month. A separate survey was developed for faculty, again focusing on the same themes although responses were too few to allow meaningful interpretation. Survey responses were read for themes within each question, and mentions of themes were then counted for each question. Overall themes regarding strengths and weaknesses were also tallied and discussed among the authors, within the context of current program capabilities, to identify challenges and opportunities for the program.

## FINDINGS

### Student Experiences

Eleven current students and thirteen M2M alumni responded to the survey; current and former student responses are summarized in Table 2 and Table 3, respectively. As only one faculty member responded, these responses are not tabulated.

**Table 2.**
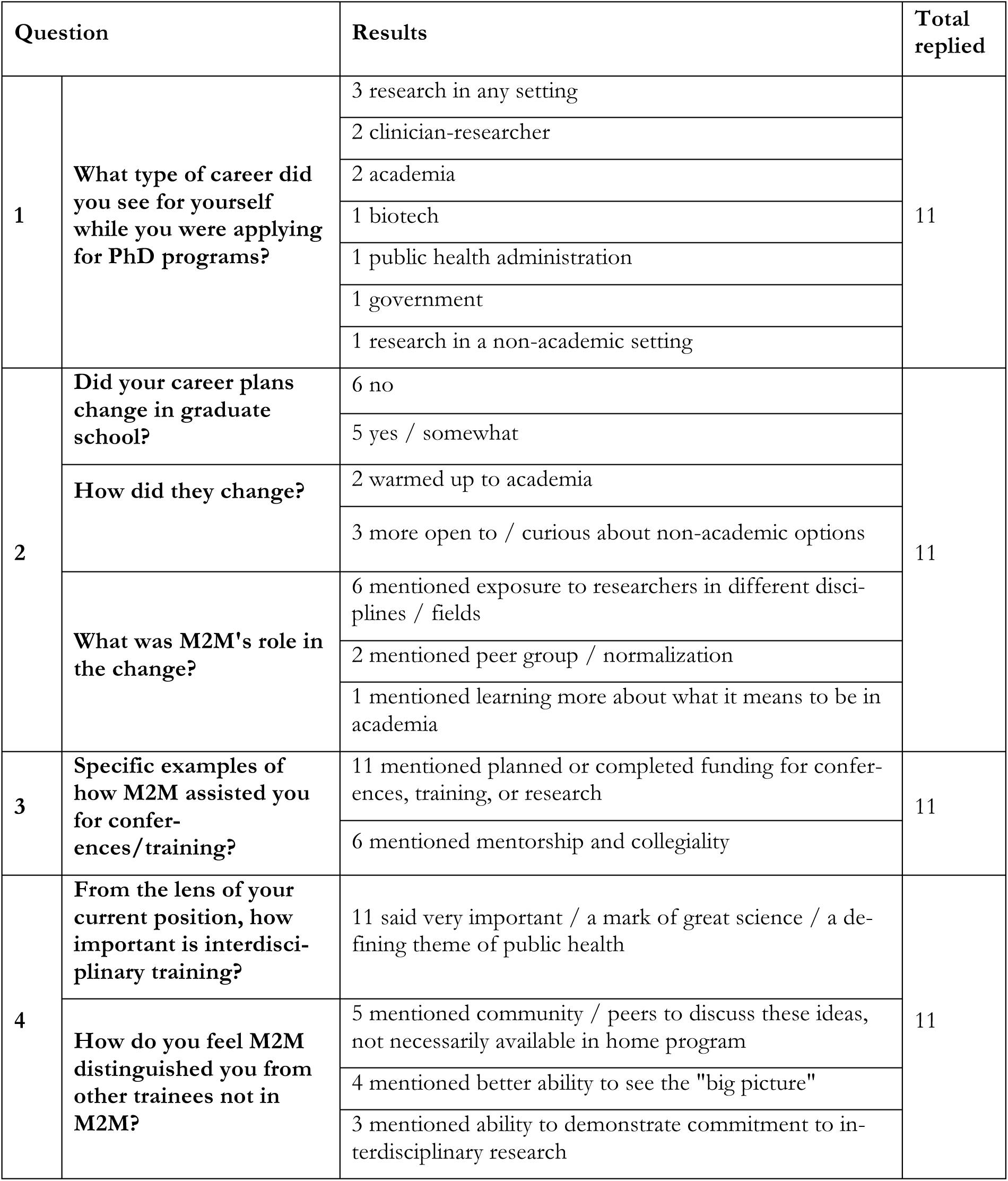
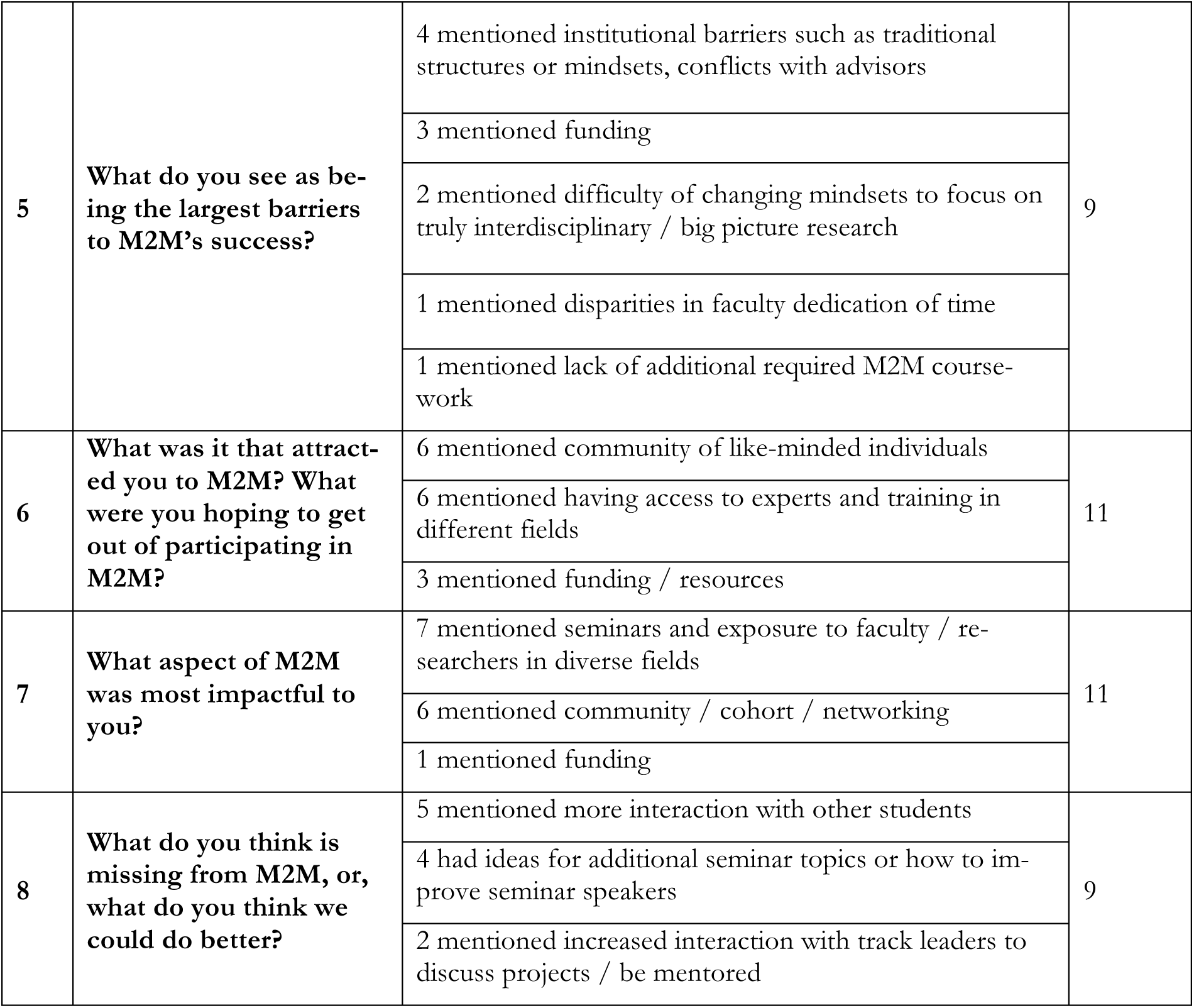
Survey responses from current M2M students, n=11. [This section is completed by the Publisher] Accepting Editor:. □ Received: (date) □ Revised: (date) □ Accepted: (date) (this is completed by Publisher. Cite as: (completed by publisher) (CC BY-NC 4.0) This article is licensed to you under a Creative Commons Attribution-NonCommercial 4.0 International <u>License</u>. When you copy and redistribute this paper in full or in part, you need to provide proper attribution to it to ensure that others can later locate this work (and to ensure that others do not accuse you of plagiarism). You may (and we encourage you to) adapt, remix, transform, and build upon the material for any non-commercial purposes. This license does not permit you to use this material for commercial purposes.

| Question |  | Results | Total replied |
| --- | --- | --- | --- |
| 1 | **What type of career did you see for yourself while you were applying for PhD programs?** | 3 research in any setting | 11 |
|  |  | 2 clinician-researcher |  |
|  |  | 2 academia |  |
|  |  | 1 biotech |  |
|  |  | 1 public health administration |  |
|  |  | 1 government |  |
|  |  | 1 research in a non-academic setting |  |
| 2 | **Did your career plans change in graduate school?** | 6 no | 11 |
|  |  | 5 yes / somewhat |  |
|  | **How did they change?** | 2 warmed up to academia |  |
|  |  | 3 more open to / curious about non-academic options |  |
|  | **What was M2M's role in the change?** | 6 mentioned exposure to researchers in different disciplines / fields |  |
|  |  | 2 mentioned peer group / normalization |  |
|  |  | 1 mentioned learning more about what it means to be in academia |  |
| 3 | **Specific examples of how M2M assisted you for conferences/training?** | 11 mentioned planned or completed funding for conferences, training, or research | 11 |
|  |  | 6 mentioned mentorship and collegiality |  |
| 4 | **From the lens of your current position, how important is interdisciplinary training?** | 11 said very important / a mark of great science / a defining theme of public health | 11 |
|  | **How do you feel M2M distinguished you from other trainees not in M2M?** | 5 mentioned community / peers to discuss these ideas, not necessarily available in home program |  |
|  |  | 4 mentioned better ability to see the "big picture" |  |
|  |  | 3 mentioned ability to demonstrate commitment to interdisciplinary research |  |
| 5 | **What do you see as being the largest barriers to M2M's success?** | 4 mentioned institutional barriers such as traditional structures or mindsets, conflicts with advisors | 9 |
|  |  | 3 mentioned funding |  |
|  |  | 2 mentioned difficulty of changing mindsets to focus on truly interdisciplinary / big picture research |  |
|  |  | 1 mentioned disparities in faculty dedication of time |  |
|  |  | 1 mentioned lack of additional required M2M coursework |  |
| 6 | **What was it that attracted you to M2M? What were you hoping to get out of participating in M2M?** | 6 mentioned community of like-minded individuals | 11 |
|  |  | 6 mentioned having access to experts and training in different fields |  |
|  |  | 3 mentioned funding / resources |  |
| 7 | **What aspect of M2M was most impactful to you?** | 7 mentioned seminars and exposure to faculty / researchers in diverse fields | 11 |
|  |  | 6 mentioned community / cohort / networking |  |
|  |  | 1 mentioned funding |  |
| 8 | **What do you think is missing from M2M, or, what do you think we could do better?** | 5 mentioned more interaction with other students | 9 |
|  |  | 4 had ideas for additional seminar topics or how to improve seminar speakers |  |
|  |  | 2 mentioned increased interaction with track leaders to discuss projects / be mentored |  |

**Table 3.** Survey responses from M2M alumni, n=13

| Question |  | Results | Total re-plied |
| --- | --- | --- | --- |
| 1 | What type of career did you see for yourself while you were applying for PhD programs? | 5 academia<br>1 academia/policy<br>1 biotech/medicine<br>2 clinical care and research<br>2 government<br>1 research unspecified<br>1 unsure | 13 |
| 2 | Did your career plans change in graduate school? | 11 yes<br>2 no | 13 |
|  | How did they change? | 5 turned away from academia<br>1 turned away from policy<br>1 turned focus to teaching<br>1 career path solidified |  |
|  | What was M2M's role in the change? | 4 mentioned “home,” community, or validation<br>9 mentioned expanded focus, interests, or opportunities |  |
| 3 | Specific examples of how M2M assisted you for conferences/training? | 11 mentioned funding for either travel or research<br>10 mentioned collaboration/diverse viewpoints, mentoring, support and collegiality<br>1 was exposed to future career | 13 |
| 4 | From the lens of your current position, how important is interdisciplinary training? | 9 said important/vital/fundamental | 13 |
|  | How do you feel M2M distinguished you from other trainees not in M2M? | 11 mentioned interdisciplinary science/collaboration/training as competitive edge<br>2 mentioned networking<br>1 mentioned not feeling distinguished because of home program pushback |  |
| 5 | **What do you see as being the largest barriers to M2M's success?** | 6 mentioned difficulty in getting students to feel comfortable in an outside field<br>5 mentioned institutional barriers such as traditional structures or mindsets, conflicts with advisors<br>5 mentioned funding | 11 |
| 6 | **What was it that attracted you to M2M? What were you hoping to get out of participating in M2M?** | 9 mentioned community of like-minded individuals<br>6 mentioned having access to train in a new field<br>3 mentioned funding | 12 |
| 7 | **What aspect of M2M was most impactful to you?** | 5 mentioned seminar class directly<br>9 mentioned the community of respect, support, mentoring, encouragement, confidence building, or open discussions<br>4 mentioned professional development, networking | 13 |
| 8 | **What do you think is missing from M2M, or, what do you think we could do better?** | 3 mentioned wanting more alumni interaction or other networking<br>3 had suggestions for seminar class topics including humanities and mental health, stress, isolation, mentor/mentee relationships<br>1 mentioned a return to funding support<br>3 basically "keep it up"<br>2 wanted more structured requirements to negotiate time with their home programs<br>1 mentioned more program outreach, within Emory to pick up struggling aspiring interdisciplinary researchers, and to the vibrant and innovative Atlanta community<br>1 mentioned wanting more time with track leaders | 12 |

When asked what type of career trainees envisioned while applying to PhD programs, the majority of responses indicated vague interest in academic health research with a few individuals interested in government research, policy, and biotechnology. Many students clearly had strong research experience at this stage and mentors that had shepherded them into pursuing their PhD, with a commonly heard trope that “a PhD was the next step” for a student studying biological science. In addressing how their career goals shifted during their PhD program, responses were divided among those who continued pursuits of academic research or teaching (often sparked by graduate school experiences in teaching assistantships), and those who pivoted away from the academic setting—for instance toward applied public health research in the government setting.

Many students reported experiencing resistance from their home program leadership and dissertation advisors against their interest in integrating population and laboratory research. All students and alumni remarked that the M2M program provided psychological support by validating their interests and pursuits, as well as their commitment to interdisciplinary science. Students stated that they felt respected by M2M program mentors more as colleagues than pupils, suggesting a possible dearth of respect within the traditional PhD training paradigm.

M2M also served as a bridge between students and their home program mentors. Students and alumni often remarked that dissertation mentors were pleased by M2M’s financial support for conferences, technical training, and small project costs. Every student was able to access funds readily that assisted them in honing career goals and gaining unique experiences through outside training opportunities or conferences. Numerous students were able to use funds to gain technical experience in external laboratories.

All student respondents indicated a belief that interdisciplinary research is absolutely the future of medical science. Most students felt emboldened by M2M and remarked on their increased confidence in discussing research across disciplines. Several students also discovered their future direction in a specific research topic or field from interactions with M2M and its invited discussion leaders. Students noted the impact of exposure to diverse fields and career paths via the weekly seminar class, believing that this exposure served an important role in broadening student perspectives.

When asked how M2M could improve, the overall theme was simply to “keep it up”—students and alumni were highly appreciative of the collegial atmosphere and ability to interact regularly with faculty and fellow students, and many requested that these opportunities be made even more frequent. M2M graduates truly feel that the program helped define their early careers and are hopeful for its continued presence in future students’ lives. Faculty have also expressed that the program approach was effective, and noted the importance of continued (and expanded) resources for students.

BWF as a funding agency has been critical to the development of M2M by encouraging ongoing adjustments to M2M’s self-defined policies. This approach has allowed the program to quickly discard requirements that became counterproductive to its training mission and to recast its design through a continuing cycle of trainee feedback to program administrators. This has allowed the program leadership to tweak the structure through time, while also enabling the funds to be stretched out in order to maximize impact.

### Opportunities and Challenges

As previously noted, M2M is a doctoral “pathway,” not a doctoral “program,” *per se*. A positive consequence of this fact is that M2M – in its present iteration – can do more to mold a student’s interests and exposures, without having to concern itself with some common features of a doctoral program such as the qualifying exam. M2M sits at the interface between more traditional home programs, providing value that does not exist in those silos. However, this outside-of-the-box approach to graduate training does come with its own set of challenges. Indeed, the largest barrier to advancing interdisciplinary training models seems to be the silos themselves. Why are there so many PhD programs? Why are there so many departments? These concepts seem so out of step with the modern processes of knowledge generation. These programmatic silos have generated significant institutional power and thus are led by individuals bent on maintaining (or rationalizing) that power. The effort required to innovate in graduate education is also a strong argument for maintaining the *status quo*. A consequence of this traditional approach is that students follow fairly prescribed paths through the program, becoming ever more deeply specialized, and are not given opportunities to distinguish themselves from others in their home program. Of course, students are distinguished from their peers by their individual dissertation research projects, but most students do not continue studying in the same subdiscipline after graduate school. Hence, students that have adequate bandwidth should be encouraged to go beyond the training provided by their home program, taking on new exposures that enable them to differentiate themselves from their peers. This represents a main motivation for students who apply to M2M.

This new interdisciplinary mentoring paradigm can lead to challenges for faculty, whose professional lives are already thinly stretched across too many commitments. One faculty member noted that time is a major barrier to engaging faculty in M2M at a level beyond mentoring of individual students. While M2M does not place requirements on its program faculty, beyond participation in the seminar class and meeting with students within their tracks, their home programs often mandate participation in teaching, recruitment activities, or committees in order to remain members of the training faculty in those home programs. Faculty may therefore be less likely to commit to a program like M2M that interfaces with not only their home program but also departments where they receive no professional benefit or reward. Decreases over time in commitment from admittedly overburdened faculty can be detrimental to the training experience for M2M students.

Another challenge for faculty has been the requirement that each student have two research mentors, one based in the laboratory sciences and one based in the population sciences. While this requirement was established to help ensure that the student’s research project successfully bridged the two fields, traditional patterns of research support for dissertation research rely upon funds from the grants of a single mentor who then retains senior authorship on the manuscripts that the dissertation research produces. Having split mentorship can complicate these understandings.

## DISCUSSION

The M2M experiment has enhanced the training of 49 PhD candidates to date, including five students engaged in the dual MD/PhD degree program. M2M’s graduates have gone on to fantastic careers as epidemiologists in federal and governmental health agencies, physician-scientists at major academic health systems, research group leaders in industry, and research faculty at large and small academic institutions, both domestic and international. Many continued in careers that included substantial ongoing research activities. All students who participated in the M2M Program consider this to be one of the highlights of their doctoral training.

Here, we highlight the words of one of the M2M alumni, Michael Mina, MD/PhD, currently Chief Resident Physician, Department of Pathology, Brigham and Women’s Hospital Harvard Medical School:

“From the start, M2M was hugely successful with both PhD candidates and their mentors, with a near frenzy among students to gain entry, and faculty willingness to participate. At first this was surprising, since faculty and PhD students rarely desire increased curricular requirements. But it owed to the ever-present interdisciplinary nature of much of the research at Emory. Despite an omnipresence of collaboration among the faculty, what was previously missing was a curriculum and a ‘pathway’ for PhD students to leverage the cross-disciplinary work of their mentors, and to be formally trained across disciplines. M2M provided that, and the famous line ‘If you build it, they will come’ couldn’t have been truer. My PhD at Emory straddled not just laboratory benchtop sciences and mathematical/population sciences, but also engaged faculty across departments, schools, and hospital settings. While I attribute much of this to the formal framework provided by the M2M program, it was rooted in a broad institutional acknowledgement of the importance of stretching across divisions to do the best science possible.”

M2M is of course not the only program to attempt to formalize training in interdisciplinary research, and the benefits that our students highlighted are similar to those highlighted in other studies: for instance, interactive teaching and smaller class sizes (e.g., in the seminar), real-world examples of interdisciplinary research questions, and frequent peer interaction and collaboration[19-21]. However, unlike many other efforts, M2M is not limited to a single course or year [19-21], enabling students to gain exposure to a breadth of topics and interdisciplinary research questions. It’s clear that programs like M2M have value: in a report commissioned by the NSF on the Integrative Graduate Education and Research Traineeship (IGERT) program, the analysis found that time to defense was shorter for students at grantee institutions, and that directors felt that the programs improved the institutional prominence of interdisciplinary research [22]. Further, it can be noted that interdisciplinary life-sciences PhDs, particularly in public health, are only increasing [23].

The task at hand for the M2M Program leadership is to find a funding stream that will enable this experimental program to continue beyond the lifetime of the BWF grant, which is non-renewable. The NIH’s T32 funding mechanism is not an obvious option, since those grants are typically geared toward funding programs that are highly specialized and clearly tied to one of the Institutes/Centers of that agency. Other potential funding agencies also do not appear to be appropriate for support of a disease-and organ system-agnostic training platform as M2M. Indeed, it appears that this experimental program that was designed to fill a major void in biomedical training is now falling into the gaps between traditional funding mechanisms for graduate training.

## CONCLUSION

Statements such as those by the alumnus above suggest that the M2M experiment has been highly successful. The challenges associated with building a training program at the interface of well-established home programs are substantial, but not insurmountable. Top trainees at an institution such as Emory seek ways to distinguish themselves during their training, to stretch themselves beyond the boundaries of the specialized training from the home program. The experience of the M2M leadership is that students who participate in M2M are, indeed, the cream-of-the-crop of trainees from each of the home programs that they represent.

The M2M Program serves as a great example of a training pathway that links major sectors of science (i.e., population-based and lab-based) by enabling students to seek training between two disciplines (e.g., genetics and epidemiology, or pharmacology and public health): this is truly interdisciplinary training. This raises the question of whether such interdisciplinary training programs lead to scientists who are jacks of all trades and masters of none. M2M avoids that concern by making sure that the genetics student in this example meets all of the requirements of the genetics PhD program, but extends herself by also acquiring substantial training in epidemiology. The end result is, at minimum, a student with a PhD in genetics who also understands the application of quantitative tools for population biology. However, the introduction of interdisciplinarity goes well beyond this, since M2M is not made up of just several genetics students seeking to understand epidemiology working with several epidemiology students seeking to understand genetics. Because each cohort of students matriculating into the program is composed of students across many disciplines, and because the program faculty also come from a wide array of backgrounds, there is added exposure across disciplines. As a consequence, students and faculty develop new ways of thinking, building bridges from the safe spaces of their fields and branching out into new intellectual paradigms that never occurred to them before.

Our hope is that by providing this description of the successes and challenges experienced by the M2M program, we may facilitate the work of other institutions in establishing programs with similar goals of enhancing interdisciplinarity in graduate education.

